# IL-15 promotes inflammatory T_h_17 cells in the intestine

**DOI:** 10.1101/2023.03.11.532227

**Authors:** Jonathan G. Golob, Guoqing Hou, Allen Lee, Helmut Grassberger, Elliott M Berinstein, Mohamed El Zataari, Valerie Khaykin, Christopher Fry, Jeff B. Berinstein, Jean Nemzek, Nobuhiko Kamada, John Y Kao, Shrinivas Bishu

**Affiliations:** Division of Infectious Diseases, University of Michigan; Division of Gastroenterology, University of Michigan; Department of Medicine, Trinity Health; Unit of Laboratory Animal Medicine, University of Michigan

## Abstract

Ulcerative Colitis (UC) is a chronic gastrointestinal condition with high morbidity. While modern medical therapies have revolutionized the care of UC, 10-25% of patients fail medications and still progress to surgery. Thus, developing new treatments is a core problem in UC. T-cells, especially T_h_17 cells, are strongly linked with UC and are major targets of medications in UC. Tissue-resident memory T-cells (T_RM_) are a distinct class of T-cells that are highly enriched in the intestine, closely aligned with the microbiota, and are implicated in the pathogenesis of UC. Unlike circulating T-cells, T_RM_ are difficult to target because they do not recirculate. Thus, we focused on cytokines like IL-15 which act as a tissue danger signal and regulate T-cells *in situ*. We found that the *IL15* axis is upregulated in UC and predicts treatment response. IL-15 was redundant for T_h_17 differentiation but could activate terminally differentiated T_h_17 cells to promote intestinal inflammation. Finally, in CD4^+^ T_RM_ from patients with UC, IL-15 upregulated *RORC*, the master transcription factor for T_h_17 cells, via a Janus Kinase (JAK)1 pathway. Thus, IL-15 promotes terminally differentiated inflammatory T_h_17 cells in the intestine raising the possibility that IL-15 may be a target for UC treatments.

## INTRODUCTION

Ulcerative colitis (UC) is an incurable, relapsing, and remitting condition that is thought to be driven by dysregulated immune responses to dysbiotic gut microbiota. UC is characterized by inflammation of the rectum, which often extends to involve the proximal colon, and presents as bloody diarrhea^1^. Modern therapies like biologics and small molecule inhibitors have substantially improved clinical outcomes in UC over the last 20 years ^1 2^. Despite their impact, it is estimated that up to 50% of patients with UC either have ‘primary non-response’ to treatments or suffer secondary ‘loss of response’ ^2^. Adding to this, flares of UC can occur unpredictably, even in patients with long-standing, well-controlled disease^1^. Thus, developing new therapies for UC remains a central challenge in Inflammatory Bowel Disease (IBD).

CD4^+^ T-cells, particularly the IL-17 producing subset of CD4^+^ T-cells (‘Th17’), are strongly implicated in the pathogenesis of UC^3^. Risk-conferring loci for UC are highly enriched for T_h_17 pathway genes and T_h_17 cells are increased in the intestine of patients with UC^4 5 6 7^. Moreover, a multitude of mouse studies demonstrates the colitogenic potential of T_h_17 cells across varied models of colitis like T-cell transfer, IL-10^-/-^, and dextran sodium sulfate (DSS)^7 8^. T_h_17 cells are defined by the production of IL-17A but it is important to recognize that IL-17A is not pathogenic in IBD. Rather, murine studies of DSS colitis indicate that IL-17A protects the intestinal epithelium, which explains the clinical finding that anti-IL-17A treatment worsens IBD outcomes^9, 10 11^. Thus, T_h_17 cells are pathogenic via IL-17A-independent mechanisms, including the production of granulocyte-macrophage colony-stimulating factor (GM-CSF, encoded by *csf2*) ^12^. Adding to this complexity, T_h_17 cells have an inherent ‘plasticity’ and can be ‘non-pathogenic’, typified by IL-10 production, or ‘pathogenic’, characterized by the production of Interferon (IFN)ɣ and GM-CSF and the expression of the IL-23 receptor (IL-23R) ^12, 13^. Thus, IL- 17A is a marker of T_h_17 cells but does not define their phenotype. T_h_17 cells are major targets of anti-IL-23 therapies in UC, underscoring their pathogenicity in IBD.

Adding another layer, T-cells are further specialized into anatomically restricted compartments^14, 15^. Peripheral tissues like the intestine are awash in a unique and distinct class of T-cells termed tissue-resident memory T-cells (T_RM_). T_RM_ can be CD8^+^ or CD4^+^, and any subtype therein (T_h_17, T_h_1), but share the core aspects that they are maintained long-term *in situ*, exhibit little recirculation, and are poised for rapid reactivation ^14–16^. T_RM_ are the most abundant intestinal T-cells in UC and are implicated in the pathogenesis of UC^17 18^. Unlike circulating T-cells which are relatively simpler to target, T_RM_ are protected from manipulation because they are disconnected from the circulation, and the challenge is to develop ways to target them. One approach is to consider the cytokines that regulate T_RM_. In this respect, IL-15 directs the homeostatic processes of committed T-cells and is highly expressed in IBD^19 20 21^. IL- 15 is required to maintain CD8^+^ T_RM_ in tissue, but the extent to which it regulates CD4^+^ T_RM_ is unclear^22^. Given all these points, we used combined *in silico* and *in vitro* approaches in patients with UC as well as murine models, to examine whether IL-15 regulates CD4^+^ T_RM_ in UC. We confirm that the IL-15 axis is upregulated in UC and find that T_h_17 T_RM_ that are associated with UC express the IL-15 receptor β-subunit (hereafter called IL-2RB because it is also a part of the IL-2R complex). Functionally, IL-15 was redundant for T_h_17 differentiation and the long-term tissue residency of intestinal T_h_17 T_RM_. However, it acted directly on T_h_17 T_RM_ to drive the production of inflammatory cytokines via Janus Kinase (JAK)1, and likely also via *RORC*, which encodes the master transcription factor for T_h_17 cells retinoic acid receptor-related orphan receptor (ROR)ɣt. Thus, our data suggest that IL-15 may drive pathogenic T_h_17 T_RM_ in UC, raising the possibility that IL-15 or IL-2RB blockade may be treatment options for UC.

## RESULTS

### The IL-15 axis is upregulated in Inflammatory Bowel Disease

IL-15, along with IL-2 and IL-7, is a member of the common gamma chain (γ_c_) family so-called because they exhibit overlapping receptor complexes^19^. This family is so fundamental to immune system homeostasis that impairment in them leads to severe combined immune deficiencies (SCID) in humans^19^. The IL-15R complex is composed of the γ_c_ subunit, which is common to the γ_c_ cytokines, and the IL-2RB subunit which confers cytokine-specific signaling^23 24^. Most biologically relevant IL-15 signaling *in vivo* is via *trans*-presentation wherein antigen-presenting cells (APC) present IL-15 in the context of IL-15RA to responder T-cells (**Fig 1A**)^24 22, 23^. The IL-2RB subunit is also a component of the IL-2R complex and is more commonly called IL-2RB, so to avoid confusion we will refer to IL-15RB as IL-2RB. The γ_c_ subunit and IL-2RB subunits signal to the downstream effectors Janus Kinase (JAK) 3 and JAK1, respectively. Of these pathways, JAK 1 and JAK3 activate STAT5, which can induce FOXP3 the master transcription factor of T-regulatory (T_reg_) cells^25 26^. By comparison, the IL-2RB-JAK1-STAT3 pathway is less well characterized (**Fig 1A**).

**Figure 1.**
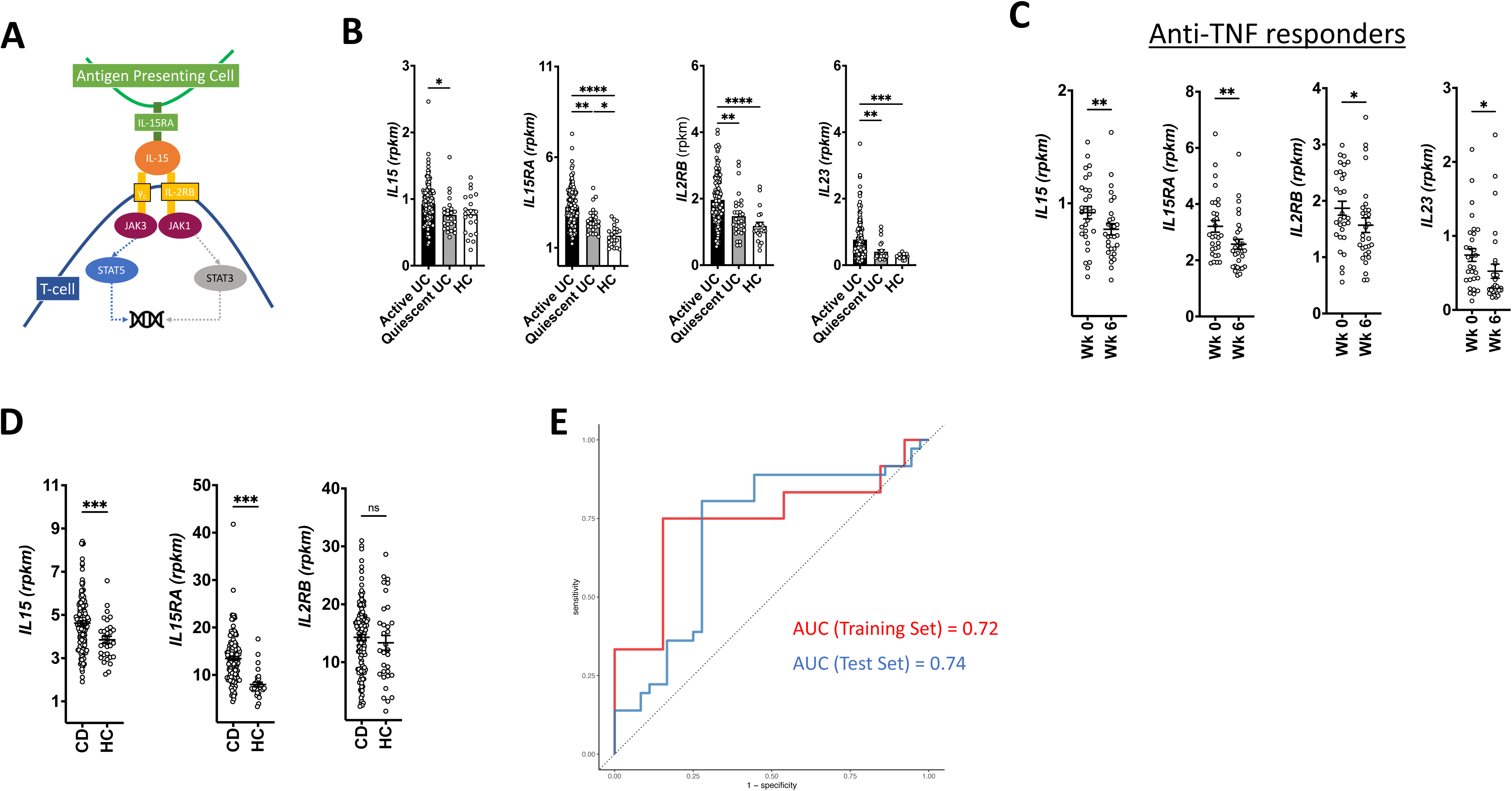
The IL-15 axis is upregulated in Inflammatory Bowel Disease. **A)** Schematic of IL-15 signaling. **B)** induction of *IL15* pathways and *IL23* from colonic specimens of patients with UC enrolled in the Golimumab clinical trial for UC. Healthy controls (HC) are also from that study. These data present patients with active (complete Mayo score(cMS) ≥ 4) and quiescent UC (cMS < 4) at any point in the trial. **C)** Changes in week 0 and 6 in responders to Golimumab. **D)** Induction of the *IL15* axis in patients with active colonic Crohn’s Disease (CD) from the pediatric IBD (RISK) cohort. E) Area under the curve (AUC) from a training and test set based on the trial data to assess the extent to which *IL15* axis transcripts predict response to Golimumab. **** to * = p < 0.0001 to p < 0.1. A responder in (C) was defined based on the trial metric.

To understand the biological relevance of the IL-15 axis in IBD, we examined the expression of *IL15*, *IL15RA,* and *IL2RB* (*IL15RB*) in UC and Crohn’s Disease (CD) using public data from the anti-TNF Golimumab clinical trial and the RISK cohort, respectively^27, 28^. Consistent with a potential role, *IL15*, *IL15RA* and *IL2RB* are all upregulated in active, relative to quiescent UC and non-IBD healthy controls (HC) (**Fig 1B**). *IL23*, which is thought to be causally implicated in UC, is presented as an additional control. Consistent with the association of the IL-15 axis with inflammation, these transcripts all decline in Golimumab responders but not in non-responders (**Fig 1C** and **Sup Fig 1A**). In CD, *IL15* and *IL15RA* are upregulated relative to controls whereas *IL2RB* was not (**Fig 1D**). To further investigate the clinical relevance of the IL-15 axis in IBD, we trained a random forest model on the *IL15* transcript levels to predict response to Golimumab in UC. The model showed good accuracy in predicting response in both the training set (Area Under the Curve (AUC) = 0.72) as well as the independent hold-out test set (AUC = 0.74) (**Fig 1E**). The most important predictors for the model by variable importance scores included *TNF*, *IL23a*, *ITGB7*, *ITGA4*, and *IL2RB* (**Sup Fig 1B**). Finally, we verified that *IL15*, *IL15RA,* and *IL2RB* were upregulated in UC by querying an independently reported meta-analysis comprising 85 mucosal gene sets in UC (last accessed March 7^th^, 2023 at https://premedibd.com/genes.html) (**Sup Fig 1C**) ^29^. Collectively, these data indicate that the IL-15 axis may be biologically relevant in IBD.

### *IL2RB* expressing CD4^+^ T_RM_ exhibit an inflammatory T_h_17 transcriptional signature

IL-15 regulates the homeostatic processes of terminally differentiated T-cells, and the most abundant terminally differentiated T-cell subsets in the human intestine are CD4^+^ and CD8^+^ T_RM_. IL-15 is required to maintain CD8^+^ T_RM_ in the intestine, but the extent to which IL-15 regulates CD4^+^ T_RM_ is less well known ^30 31, 32^. Thus, we focused on *IL2RB* expressing CD4^+^ and CD8^+^ T_RM_ in the intestine in human UC.

As a first step, we performed an *in silico* analysis of *IL2RB* (*IL15RB*) expressing intestinal T_RM_ from publicly available single-cell sequencing (scRNA-seq) data. We identified intestinal T_RM_ as CD45RO^+^CCR7^-^CD69^+^ cells within the CD8^+^ and CD4^+^ fractions. CD103 is a commonly used T_RM_ marker. However, many CD4^+^ T_RM_ are CD103^-^ and the CD8^+^ group defined this way encompasses most CD103^+^ CD8^+^ T_RM_^14, 15 16, 33 34 35^. Thus, we did not use CD103 as a T_RM_ marker in our analysis to ensure that we captured CD103^-^ CD4^+^ T_RM_. Quantifying this data revealed an increase in the fraction of *IL2RB* expressing CD4^+^ T_RM_ in UC relative to controls (**Table 1**). In contrast, the fraction of *IL2RB* expressing CD8^+^ T_RM_ declined in both UC and CD relative to controls.

**Table 1.**
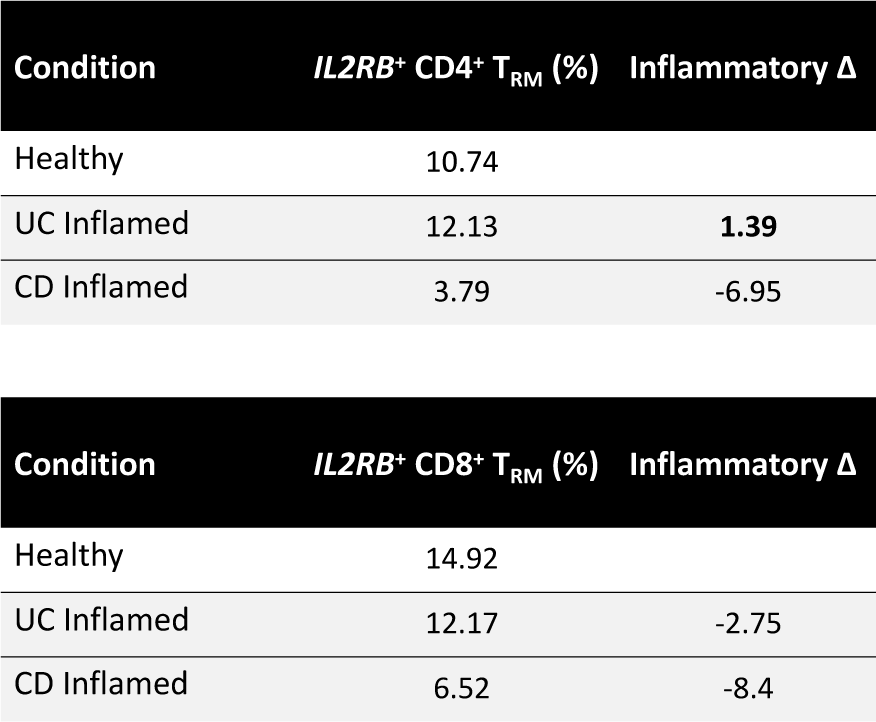

To better understand the transcriptional signature of intestinal *IL2RB* expressing T_RM_, we next assessed disease pathways and genes in *IL2RB* expressing T_RM_. Intriguingly, the top term in *IL2RB* expressing CD4^+^ T_RM_ is IBD (**Table 2**). Moreover, the genes (*IL23R*, *IL17A MAF*) most strongly associated with *IL2RB* CD4^+^ T_RM_ are T_h_17 signature genes. *MAF* encodes a transcription factor that upregulates the core T_h_17 genes *Rorc*, *Il17a*, and *Il23r* in murine models ^12, 13^ . There were also notable associations with immune-checkpoint genes (*TNFRSF9*, *TNFRSF4*, *CD70*, *PDCD1*, *CTLA4*) typical of T_RM_ ^15^(**Table 2**). Interestingly, the top differentially expressed genes in I*L2RB*^+^ vs *IL2RB*^-^ CD4^+^ T_RM_ are *IL32* and *CCL5*. Both of these are pro-inflammatory and chemotactic, and *IL32* and stimulates the production of inflammatory cytokines including tumor necrosis factor (TNF)⍺, IL-6, and IL-8, all of which are abundant in active UC^36, 37^ (**Table 3**). In contrast, intestinal *IL2RB* expressing CD8^+^ T_RM_ were not associated with IBD or gastrointestinal disease conditions, except non-alcoholic fatty liver disease (full gene list is provided in the methods) (**Table 4**). These data indicate that *IL2RB* expressing CD4^+^ T_RM_ may be inflammatory T_h_17 T_RM_.

**Table 2.** *IL2RB* CD4+ T_RM_

| Term | Adjusted P-value | Odds Ratio | Combined Score | Genes |
| --- | --- | --- | --- | --- |
| Inflammatory bowel disease | 0.02 | 15.88 | 137.19 | MAF;IFNG;IL23R;IL17A |
| Cytokine-cytokine receptor interaction | 0.02 | 6.04 | 49.37 | IFNG;CCL20;CD70;IL23R;TNFRSF9;TNFRSF4;IL17A |
| Rheumatoid arthritis | 0.03 | 10.87 | 79.01 | IFNG;CCL20;CTLA4;IL17A |
| T cell receptor signaling pathway | 0.03 | 9.67 | 66.24 | IFNG;CTLA4;PDCD1;CD3D |
| Th17 cell differentiation | 0.03 | 9.38 | 63.31 | IFNG;IL23R;CD3D;IL17A |
| PD-L1 expression and PD-1 checkpoint pathway in cancer | 0.14 | 8.33 | 41.74 | IFNG;PDCD1;CD3D |
| Th1 and Th2 cell differentiation | 0.14 | 8.05 | 39.59 | MAF;IFNG;CD3D |
| IL-17 signaling pathway | 0.14 | 7.87 | 38.25 | IFNG;CCL20;IL17A |
| Allograft rejection | 0.18 | 13.15 | 58.57 | IFNG;HLA-E |
| Graft-versus-host disease | 0.19 | 11.83 | 50.43 | IFNG;HLA-E |

**Table 3.** DEGs by *IL2RB* CD4^+^ T_RM_

|  | coef | q |
| --- | --- | --- |
| IL32 | 0.42466294 | 0.00031007 |
| CCL5 | 0.39692017 | 0.00053295 |
| TFEC | -29.2321 | 0.00918771 |
| DOCK4 | -17.542394 | 0.01521124 |
| MGST1 | -19.26141 | 0.01592225 |
| IGHA2 | -0.4492494 | 0.02271952 |
| CLEC11A | -12.586075 | 0.02287661 |
| MPEG1 | -19.005124 | 0.02295341 |
| FPR3 | -28.592903 | 0.03868099 |
| CTSW | 0.28572914 | 0.09583549 |

**Table 4.** *IL2RB* CD8^+^ T_RM_

| Term | Adjusted P-value | Odds Ratio | Combined Score |
| --- | --- | --- | --- |
| Parkinson disease | 1.07E-18 | 7.37 | 346.19 |
| Prion disease | 2.09E-17 | 6.59 | 285.53 |
| Amyotrophic lateral sclerosis | 1.06E-15 | 5.24 | 204.38 |
| Huntington disease | 2.76E-14 | 5.43 | 192.26 |
| Alzheimer disease | 1.40E-13 | 4.76 | 159.83 |
| Pathways of neurodegeneration | 3.74E-12 | 3.96 | 119.26 |
| Non-alcoholic fatty liver disease | 2.23E-11 | 6.93 | 195.23 |
| Oxidative phosphorylation | 1.08E-08 | 6.39 | 139.81 |
| Thermogenesis | 1.62E-07 | 4.32 | 82.27 |
| Proteasome | 1.80E-06 | 10.57 | 174.73 |

### IL-2RB is expressed by terminally differentiated inflammatory T_h_17 cells

These data raise the possibility that inflammatory T_h_17 T_RM_ in UC can respond to IL-15. Thus, we first considered whether IL-15 can direct the differentiation of T_h_17 cells. Prior work *in vitro* with exogenous IL-15 or T-cells from IL-15^-/-^ mice indicates that IL-15 restrains T_h_17 cell differentiation and favors the generation of FOXP3^+^ T_reg_ . However, biologically relevant IL- 15 signaling *in vivo* is via IL-15RA *trans*-presentation, meaning that IL-15 alone may have different biological properties than IL-15/IL-15RA ^24 23^. Additionally, T-cells from IL-15^-/-^ mice are predisposed towards T_h_17 differentiation at baseline compared to wild-type T-cells ^25 26^. It is therefore possible that the published experimental conditions are skewed toward the conclusion that IL-15 restrains T_h_17 cell differentiation.

To circumvent these possibilities, we performed T_h_17 differentiation assays using wild-type naïve CD4^+^ T-cells with IL-15/IL-15RA complex. IL-15 presented in the context of IL-15RA substantially enhances the binding affinity of responder T-cells for IL-15 and is more biologically relevant because it models *in vivo* IL-15/IL-15RA *trans*-presentation. Using this system, we found that IL-15/IL-15RA did not enhance the differentiation of naïve T-cells into T_h_17 cells in mice or humans, indicating that IL-15 is dispensable for T_h_17 differentiation (**Fig 2A, B**).

**Figure 2.**
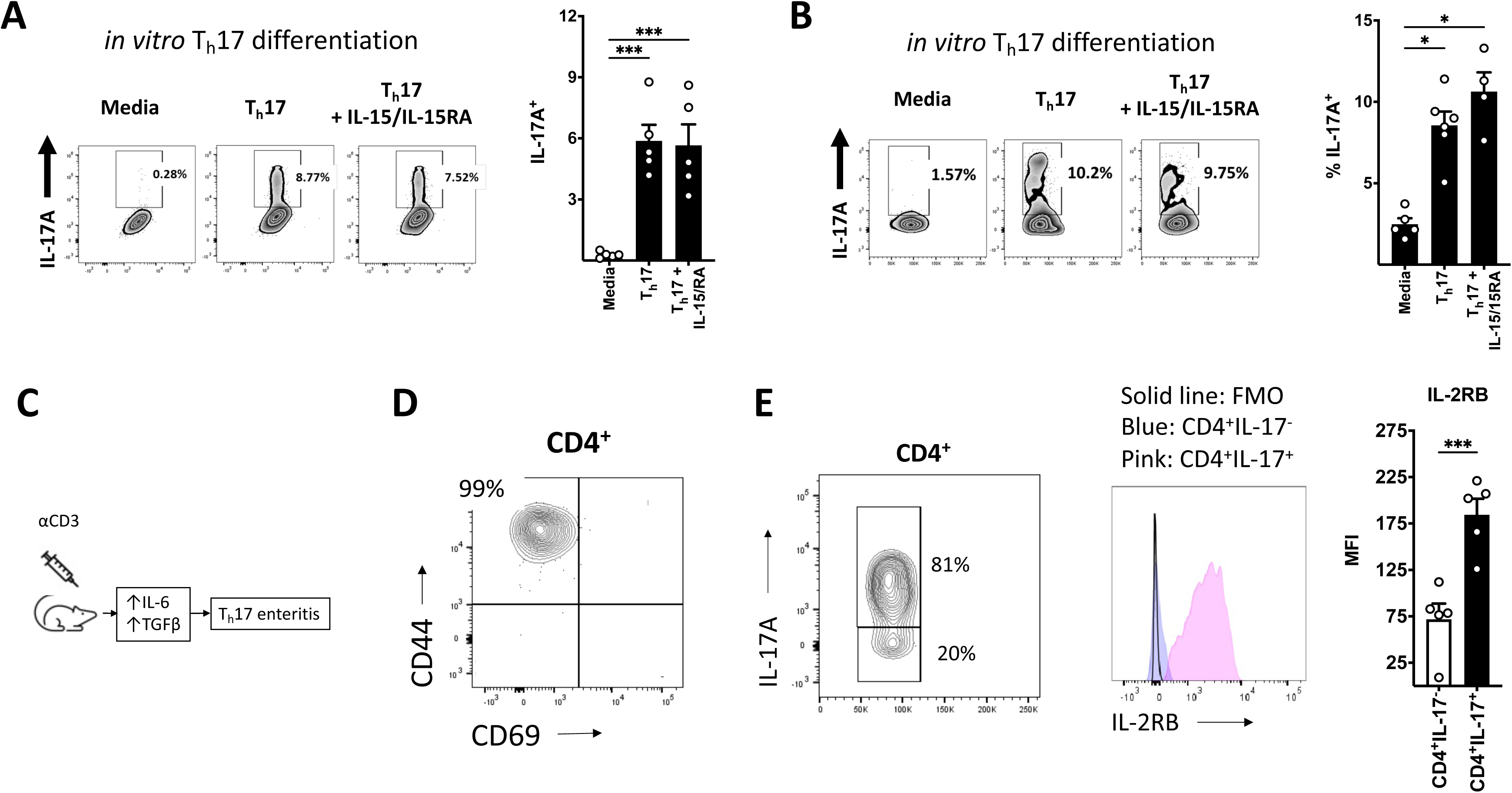
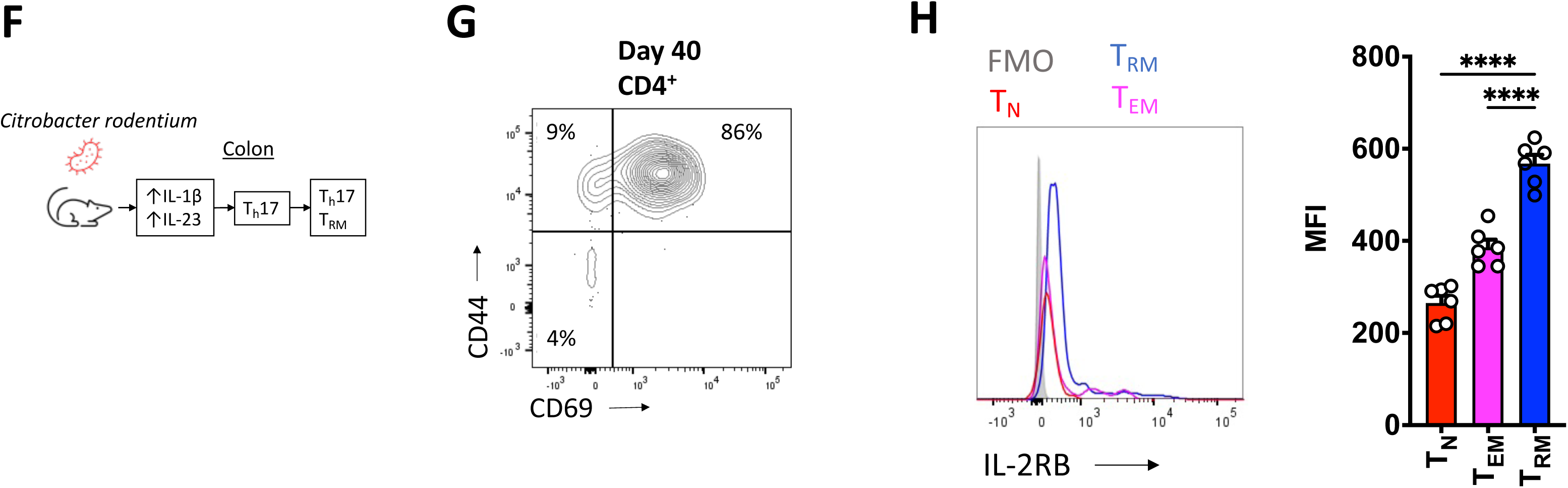
IL-2RB is expressed by terminally differentiated inflammatory T_h_17 cells. **A, B)** Naive murine **(A)** and human **(B)** CD4^+^ T-cells were *in vitro* polarized to T_h_17 cells with or without murine and human IL-15/IL-15RA complex (respectively) (left panels), and data were quantitated (right panels). **C)** Schematic of anti-CD3 enteritis, with **(D)** gating on responder CD4^+^ T-cells verifying they are T_EM_ (CD44^+^CD69^-^)**. E)** The CD4^+^ T_EM_ compartment is enriched for T_h_17 cells (left panel) which preferentially express IL-2RB relative to non-T_h_17 CD4^+^ T_EM_ (center and right panels). **F)** Schematic of *C. rodentium* induced T_h_17 cells with **(G)** gating on induced CD4^+^ T-cells at day 40 post-infection verifying most are T_RM_ (CD44^+^CD69^+^). **H)** The relative expression of IL-2RB (left panel) on naive T-cells (T_n_; CD44^-^CD69^-^), T_EM_ and T_RM_ post *C. rodentium* infection with quantified data (right panel). **** to * = p < 0.0001 to p < 0.1. Abbreviations: FMO (fluorescence minus one), MFI (mean fluorescence intensity), TGFβ (tumor growth factor β).

The differentiation of naïve CD4^+^ T-cells into committed lineages is distinct from the secondary activation of terminally differentiated T-cells. This distinction is often overlooked but is especially important for T_h_17 cells which exhibit a complex sequence of terminal differentiation and plasticity ^12, 13^. For example, IL-23 is redundant for the differentiation of naïve T-cells into T_h_17 cells, but it is required to maintain terminally differentiated T_h_17 cells. Accordingly, the IL-23R is expressed by committed T_h_17 cells, but not by naïve T-cells. Thus, we considered the possibility that IL-15 may act on terminally differentiated T_h_17 cells. To examine whether IL-2RB is expressed by terminally differentiated inflammatory T_h_17 cells in the intestine, we utilized models that generate inflammatory T_h_17 T-effector memory (T_EM_) and T_RM_ cells, respectively. T_EM_ and T_RM_ are distinct subtypes of terminally differentiated T-cells with distinct transcriptional and functional profiles and are the most common subtypes of terminally differentiated CD4^+^ T-cells in the intestine.

Intraperitoneal injection of anti-CD3 induces inflammatory T_h_17 cells in the small intestine which cause enteritis^38^ (**Fig 2C**). As we had anticipated, responder CD4^+^ T-cells after anti-CD3 injection exhibit a T_EM_ T_h_17 phenotype, which is consistent with the acute nature of the model (**Fig 2D**). Anti-CD3 enteritis can be ameliorated by blocking the trafficking of these responder (T_EM_) T_h_17 cells into the intestine^38^. Consistent with IL-15 promoting inflammatory T_h_17 cells, T_h_17 cells in anti-CD3 enteritis specifically expressed IL-2RB compared to non-T_h_17 cells (**Fig 2E**). Next, to examine the expression of IL-2RB on inflammatory T_h_17 T_RM_, we used the *Citrobacter rodentium* murine model. *C. rodentium* is a natural murine pathogen that causes a self-limiting gastrointestinal infection. *C. rodentium* specifically induces robust T_h_17 responses which transition into T_h_17 T_RM_ ^34 35 33^(**Fig 2F**). *C. rodentium* induced T_h_17 T_RM_ exhibit an inflammatory profile relative to homeostatic T_h_17 T_RM_. After clearing *C. rodentium*, most colonic CD4^+^ T-cells are T_RM_, and consistent with the scRNA seq data (Fig 2), they expressed the highest levels of IL-2RB relative to other T-cell compartments (**Fig 2G, H**). These data indicate that while IL-15 is dispensable for the differentiation of T_h_17, it may act on terminally differentiated T_h_17 cells.

### IL-15 is not required for the maintenance of pathogen-induced T_h_17 CD4^+^ T_RM_

Our data show that IL-2RB is expressed by terminally differentiated T_h_17 CD4^+^ T_RM_ in the intestine, raising the possibility that IL-15 may act on committed T-cells. The IL-15R complex on responder T-cells is composed of the γ_c_ and the IL-2RB subunits. Nearly all T-cells express the γ_c_ meaning any T-cells expressing IL-2RB have the capacity for IL-15 signaling. Functionally, IL-15 regulates the homeostatic proliferation of terminally differentiated T- and NK-cells, the migration of γδ T-cells, and the tissue residency of CD8^+^ T_RM_. We, therefore, considered whether IL-15 may analogously regulate the retention of CD4^+^ T_RM_ in tissue.

To test this, we used *C. rodentium* which induces T_h_17 (CD4^+^) T_RM_. While the optimal way to track T_RM_ is with antigen-specific transgenic T-cell receptor systems, these systems do not exist for *C. rodentium* induced T_h_17 T_RM_. Investigators circumvent this limitation by using the facts that *C. rodentium* specifically induces T_h_17 cells and that CD69 is a T_RM_ marker. Thus, it has been shown that CD44^+^CD69^+^CD4^+^ T-cells in the intestine of mice at late time points after clearing *C. rodentium* are predominantly T_h_17 T_RM_^33, 34 35^.

We coupled IL-15^-/-^ mice with this model to assess the role of IL-15 on T_h_17 T_RM_. At baseline, IL-15^-/-^ mice did not exhibit low fractions of CD4^+^ or T_h_17 cells (**Sup Fig 2A**). IL-15^-/-^ mice had very low numbers of CD8^+^ and CD3^+^CD8^-^CD4^-^ cells consistent with the well-known role of IL-15 in CD8^+^ and NK cell biology (**Sup Fig 2B**). Immunity against *C. rodentium* is strongly dependent on T_h_17 cells and independent of CD8^+^ or NK cells^39^. IL-15^-/-^ mice had an early but transient impairment in host defense to *C. rodentium* with higher bacterial counts and more weight loss (**Fig 3A**). However, IL-15^-/-^ mice did not exhibit defects in total CD4^+^ or T_h_17 cells with *C. rodentium* (**Sup Fig 2C, Fig 3B**). The T-cells in these *ex vivo* assays were stimulated with PMA and Ionomycin (P/I), a supraphysiologic stimulus typically used for flow cytometry (FCS) data. To explain the discrepancy between impaired T_h_17 function *in vivo* (host defense) and the *ex vivo* FCS data, we considered the possibility that supraphysiologic P/I stimulation can bypass any impairment due to IL-15 deficiency. Consistent with this possibility, IL-17A was very low in colon explants from IL-15^-/-^ mice compared to controls (**Fig 3C**). But mucosal IL-23, *il15ra,* and *il2rb* were normal, implying that the low IL-17A was not a consequence of low T_h_17 activating cytokines or reduced IL-2Rb expression in IL-15^-/-^ mice (**Sup Fig 2D, E**). Finally, IL-15^-/-^ mice had normal fractions and numbers of total CD4^+^ T_RM_ and T_h_17 T_RM_ at late times after clearing *C. rodentium* (**Fig 2D, E**). These data strongly imply that IL-15 is not required to maintain *C. rodentium* induced T_h_17 (CD4^+^) T_RM_. However, the lower intestinal IL-17A, which is largely produced by T_h_17 cells at these time points during *C. rodentium* infection, raise the possibility that IL-15 may impact the function of T_h_17 CD4^+^ T_RM_^33, 39^.

**Figure 3.**
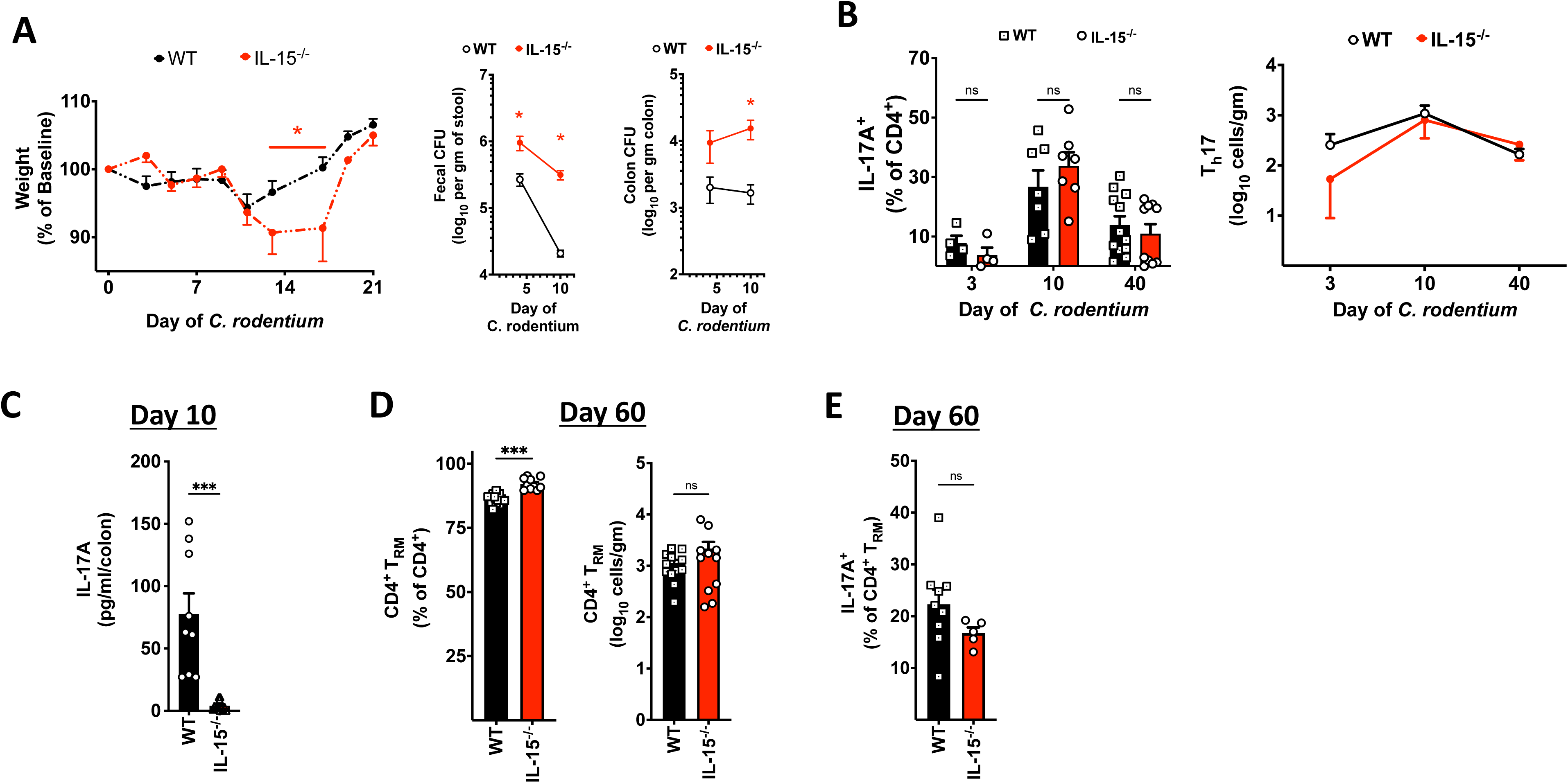
IL-15 is not required for the maintenance of pathogen-induced T_h_17 CD4^+^ T_RM_. **A)** Weight change (left panel) and bacterial burden in the stool and colon mucosa (right panels) in mice with *C. rodentium* infection. **B)** The fraction (left panel) and total number (right panel) of T_h_17 cells in mice with *C. rodentium* infection. **C)** IL-17 protein from colon explants at day 10 of *C. rodentium* infection. **D)** The fraction and the total number of CD4^+^ T_RM_ (CD44^+^CD69^+^) and **(E)** the fraction of T_h_17 T_RM_ (IL-17A^+^ CD4^+^ T_RM_) in the colon of mice at day 60 post *C. rodentium* infection. Experiment (A) is representative of 2 experiments with 5 mice per group. *** to * = p < 0.001 to p < 0.1.

### IL-15 deficient mice are protected from chemical colitis and T_h_17-driven enteritis

To assess the possibility that IL-15 may drive colitogenic T_h_17 T_RM_ *in vivo*, we coupled *C. rodentium* infection to generate T_h_17 T_RM_, followed by dextran sodium sulfate (DSS) (**Fig 4A**). While DSS is considered an ‘innate’ model because RAG^-/-^ mice develop colitis, DSS can be used to study the contribution of T-cells since T-cells can modulate the severity of DSS-induced injury^40^. We favored this approach over T-cell transfer (TCT) colitis (into RAG^-/-^IL-15^-/-^ double knockout mice). This is because TCT does not generate T_RM_, though it is a direct T-cell model of colitis. Moreover, T_RM_ cannot be transferred to recipient mice because colonic T_RM_ will not home to the colon, precluding TCT of donor T_h_17 T_RM_. Thus, while DSS may not be a direct T-cell colitis model, we opted for this approach due to the limitations of T_RM_ biology.

**Figure 4.**
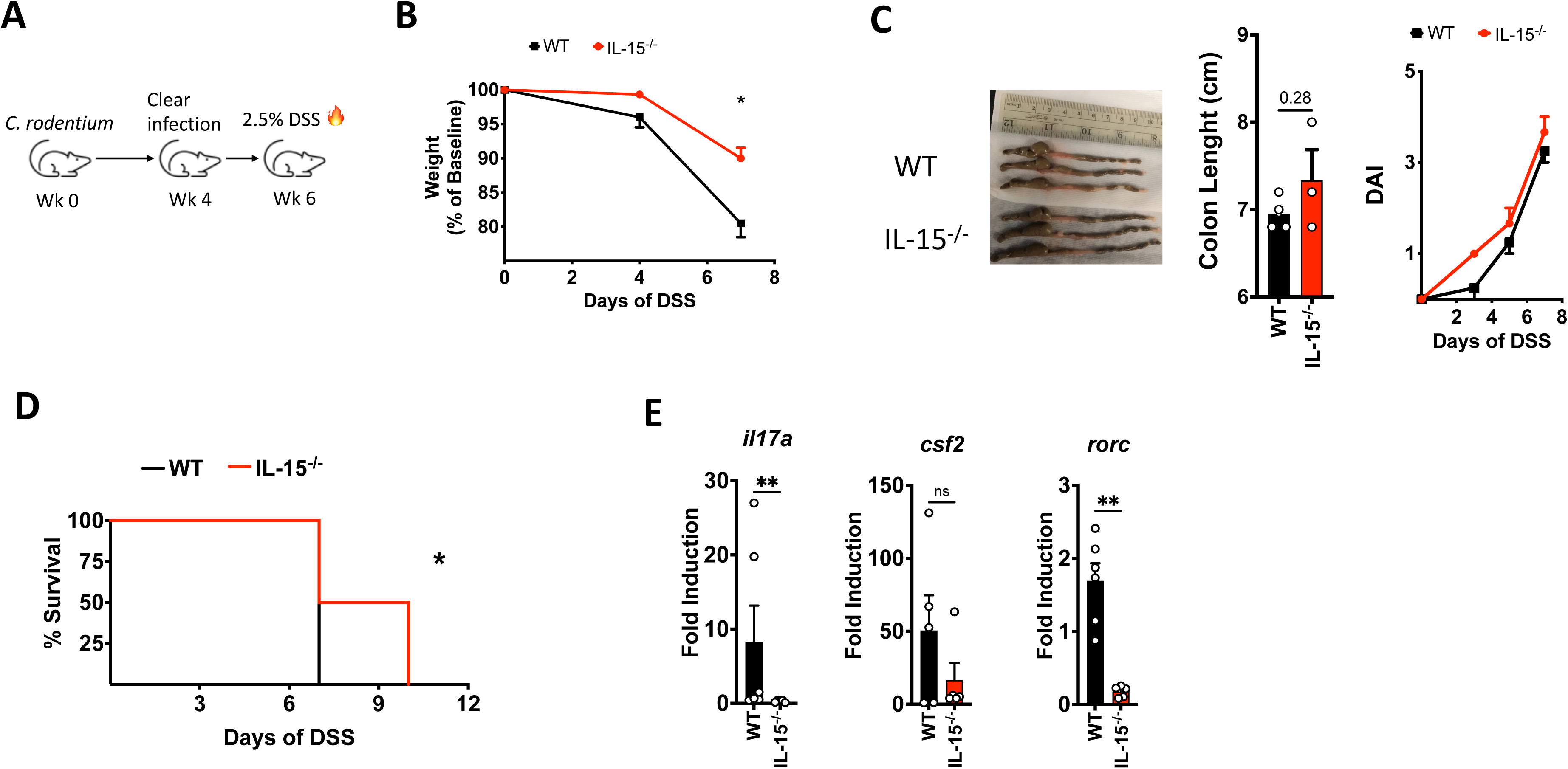
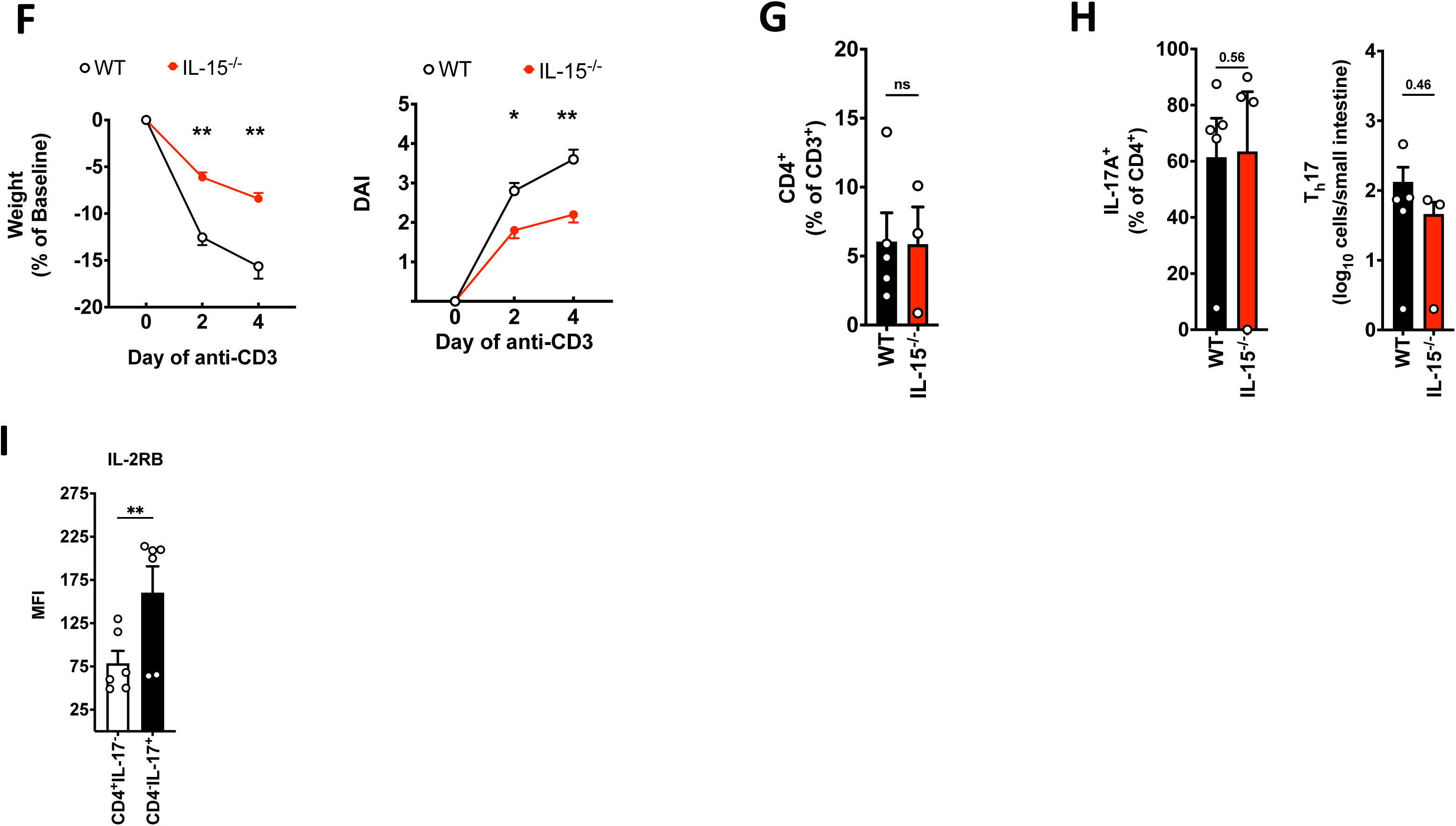
IL-15 deficiency protects against chemical colitis and T_h_17 enteritis. **A)** Experimental schematic. **B)** Weight change, **(C)** colon length (left panels) and DAI score (right panel) and **(D)** survival of mice subject to DSS as in **(A)**. **E)** Induction of the indicated genes from the colon of mice at day 7 of DSS as in **(A)**. **F)** weight change (left panel) and DAI (right panel) of mice subject to anti-CD3 enteritis as in Fig 3C. The fraction of **G)** CD4^+^ T-cells and the **(H)** fraction (left panel) and total T_h_17 cells (right panel) in the small intestine of mice at day 4 of anti-CD3 enteritis. **I)** Expression of IL-2RB on responder T_h_17 cells in the small intestine of IL-15^-/-^ mice at day 4 of anti-CD3 enteritis. All data are representative of 2 experiments with 3-5 mice per group, per experiment. ** to * = P<0.01 to p <0.1.

IL-15^-/-^ exhibited less weight loss with DSS, though colon length and disease activity were similar relative to control mice (**Fig 4B, C**). Most importantly, IL-15^-/-^ mice exhibited longer survival than controls (**Fig 4D**). Consistent with IL-15 signaling promoting pro-inflammatory T_h_17 T_RM_, *il17a*, *rorc,* and *csf2*, are significantly lower in the intestine of IL-15^-/-^ mice (**Fig 4E**). Since the IL-15R complex is also expressed on T_EM_, we also assessed the impact of IL-15 using anti-CD3 enteritis, which is driven by inflammatory T_h_17 T_EM_ (**Fig 2C-E**). As anticipated, IL-15^-/-^ mice were protected from enteritis, even though they had no defects in CD4^+^ or T_h_17 compartments in the small intestine (**Fig 4F-H**). Moreover, IL-15^-/-^ mice exhibited no defects in the expression of IL-2RB on responder T_h_17 cells (**Fig 4I vs 2E**). Thus, these data are consistent with the *in silico* and *in vitro* data indicating that IL-15 can act on inflammatory T_h_17 T_RM_ in UC.

### IL-15 promotes inflammatory T_h_17 T_RM_ in UC via Janus Kinase 1 dependent pathways

Our data suggest that IL-15 may act on inflammatory T_h_17 T_RM_ but is redundant for T_h_17 differentiation and the maintenance of T_h_17 T_RM_ *in situ*. Intestinal CD4^+^ T_RM_ from mice that have cleared *C. rodentium* is enriched for *Cr*-specific T_h_17 T_RM_. Moreover, these T_h_17 T_RM_ are considered to have an inflammatory signature relative to homeostatic CD4^+^ T_RM_.

To examine whether IL-15 acts on inflammatory T_h_17 CD4^+^ T_RM_, we used flow cytometry-assisted sorting (FACS) to purify CD4^+^ T_RM_ from the intestine of mice that had cleared *C. rodentium* (‘*Cr* CD4^+^ T_RM_’) (Gating **Sup Fig 3A**). As a biological negative control, we purified homeostatic CD4^+^ T_RM_ from uninfected age- and gender-matched mice maintained in SPF conditions (‘*Hm* CD4^+^ T_RM_’) (**Fig 5A**). Given the possibility that P/I can bypass IL-15-driven effects, we stimulated these cells with anti-CD3/CD28 and IL-15/IL-15RA complex to simulate *in vivo* T-cell receptor (TCR) ligation and trans-presentation^12, 41^. As positive controls, we also stimulated cells with the prototypically inflammatory T_h_17 cytokines IL-1β/IL-23 (with TCR ligation), or P/I. IL-15/IL-15RA complex upregulated *il17a* and *csf2* (which encodes GM-CSF), in *Cr* CD4^+^ T_RM_ nearly 2-fold compared to its effects on *Hm* CD4^+^ T_RM_ (**Fig 5B**). IL-1β/IL-23 and P/I upregulated *il17a* and *csf2*, with P/I leading to the most robust induction consistent with its supraphysiologic stimulation. In contrast, induction of *il23r* was restricted to IL-1β/IL-23, which is consistent with the literature, and *il2* was induced under all conditions verifying that cells were viable (**Sup Fig 3C**).

**Figure 5.**
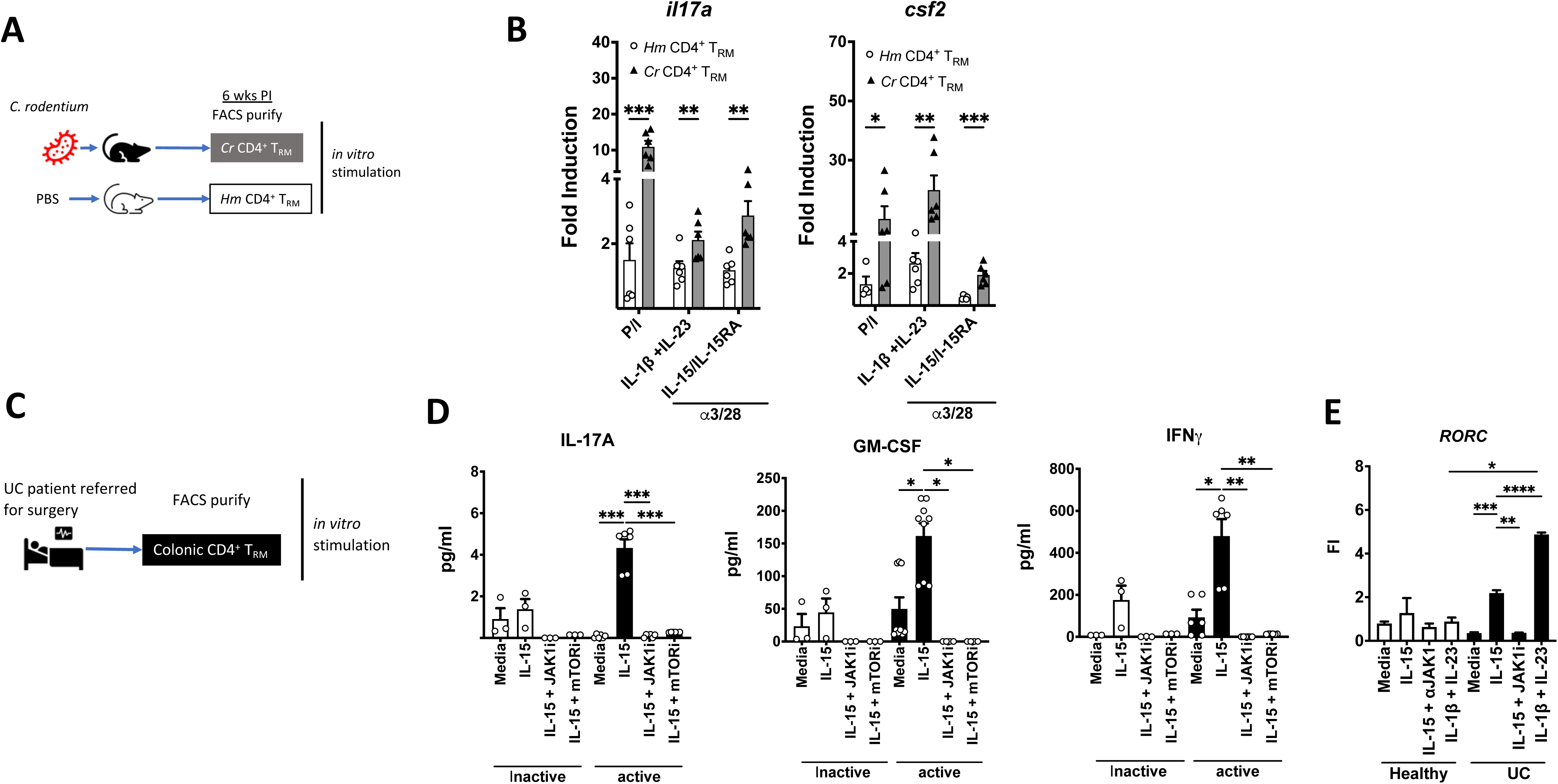
IL-15 promotes inflammatory T_h_17 T_RM_ in UC via Janus Kinase 1 dependent pathways. **A)** Homeostatic (*Hm*) and *C. rodentium* (*Cr*) induced CD4^+^ T_RM_ were FACS purified from the colon of mice and **(B)** the induction of *il17a* and *csf2* was determined after cells were stimulated as indicated. **C)** CD4^+^ T_RM_ were FACS purified from active and inactive regions from the colon of patients with medically refractory UC. As indicated, the purified CD4+ TRM were stimulated with anti-CD3 and CD28 with or without cytokines and inhibitors, and the production of **D)** IL-17A, GM-CSF, and interferon γ and the **(E)** induction of *RORC* was determined. All points are biological replicates. Genes and normalized to (B, E) GAPDH, and conditions are normalized to (B) *Hm* CD4^+^ T_RM_ and (E) media. **** to * = p < 0.0001 to p < 0.1. Murine CD4^+^ T_RM_: Live CD3^+^CD4^+^CD44^+^CD69^+^. Human CD4^+^ T_RM_: Live CD3^+^CD4^+^CD45RO^+^CD69^+^CCR7^-^.

To validate these findings in humans, we FACS purified CD4^+^ T_RM_ from colonic areas with inactive and active disease, of patients with UC referred for surgery (**Fig 5C, Gating Sup Fig 3B**). The degree of histologic activity was verified by a blinded gastrointestinal pathologist (further details are in the methods). The IL-15R complex, composed of the γc and IL-2RB subunits, signals via JAK3 and JAK1, respectively (**Fig 1A**)^19^. STAT3 regulates the transcription of both *IL17A* and *RORC* (which encodes RORɣt, the master transcription factor of T_h_17 cells) ^42, 43^. Accordingly, STAT3 is critical in T_h_17 cell biology, and humans with impaired STAT3 activation exhibit primary immune deficiencies of T_h_17 cells. Therefore, we considered the possibility that IL-15 may regulate T_h_17 T_RM_ via an IL-2RB-JAK1-RORC pathway. The purified CD4^+^ T_RM_ were stimulated via TCR ligation with a JAK1 inhibitor (**Fig 5D, E**). Because IL-15 also stimulates homeostatic proliferation by regulating the mechanistic target of rapamycin (mTOR), we also included the mTOR inhibitor rapamycin as an additional control (**Fig 5D**) ^44^. We measured the production of IL-17A, GM-CSF, and interferon (IFN)ɣ, because these cytokines are strongly associated with inflammatory T_h_17 cells (**Fig 5D**). As we anticipated, IL-15-driven production of inflammatory T_h_17 cytokines was completely blocked by inhibiting JAK1. Furthermore, IL-15/IL-15RA complex also upregulated *RORC*, though not to the same extent as IL-1β/IL-23 which is known to strongly induce *RORC*. The upregulation of *RORC* was also ameliorated by blocking JAK1.

In these assays, it is important to recognize that IL-17A is a marker of T_h_17 cells but is not itself pathogenic in IBD. However, T_h_17 cells are the targets of approved therapies in IBD and are pathogenic in IBD via IL-17A-independent mechanisms, like the production of GM-CSF. Thus, this data is consistent with the paradigm wherein IL-15/IL-15RA acts on terminally differentiated, inflammatory T_h_17 T_RM_ in UC, via an IL-2RB-JAK1-RORC pathway.

## DISCUSSION

Herein, we corroborate that the *IL15* axis is upregulated in IBD and that colonic CD4^+^ T_RM_ that express *IL2RB* (the cytokine specificity conferring subunit of the IL-15R) exhibit an inflammatory T_h_17 signature. Moreover, these *IL2RB* expressing T_h_17 T_RM_ are linked with IBD and are enriched in UC. IL-15 was redundant for the primary differentiation of murine and human T_h_17 cells, but IL-2RB was expressed on terminally differentiated T_h_17 T_EM_ and T_RM_. In contrast to the role of IL-15 on CD8^+^ T_RM_, IL-15 does not appear to be necessary to maintain CD4^+^ T_RM_ *in situ*. Instead, IL-15 acted directly on CD4^+^ T_RM_ to stimulate the production of inflammatory T_h_17 cytokines. This IL-15-driven cytokine production and upregulation of *RORC*, the master transcription factor for T_h_17 cells, and could be ameliorated by inhibiting JAK1. Finally, IL-15^-/-^ mice exhibited downregulation of T_h_17 pathways and were protected from DSS and T_h_17-driven enteritis.

Our data indicate that IL-15 is pro-inflammatory on T_h_17 T_RM_ in UC. However, IL-15 biology is complex involving *cis*- and *trans*-presentation (in NK cells, terminally differentiated CD8^+^ and CD4^+^ T-cells, and CD8^+^ T_RM_), and autocrine signaling (in antigen presenting cells (APCs) ^19, 22–24^. Prior studies on IL-15 and T_reg_/T_h_17 cell balance have used *1) in vitro* T_h_17 differentiation assays with IL-15^-/-^ mice, or *2)* IL-15 alone without IL-15RA complex, *3)* purified populations of terminally differentiated cells (specifically T_regs_), or *4)* mixtures of cells containing multiple populations of differentiated subsets (like total CD4^+^ T-cells) or *5)* have used IL-15^-/-^ deficient hosts^25, 26^. These systems are all reasonable, but not without caveats. Naïve T-cells from IL-15^-/-^ mice consistently exhibit an enhanced capacity for T_h_17 differentiation *in vitro*^25, 26^. However, *in vitro* T_h_17 differentiation assays only contain TCR ligation and cytokines, but not exogenous or APC derived IL-15. Therefore, the capacity of naïve T-cells from IL-15^-/-^ mice to have enhanced T_h_17 differentiation suggests off-target effects of IL-15 deficiency, rather than a direct impact of IL-15 on T_h_17 differentiation. Moreover, IL-15 signaling *in vivo* to responder T-cells is primarily via trans-presentation and IL-15 alone has a much lower binding affinity than IL-15/IL-15RA complex. It is therefore possible that IL-15 has differing biological functions than IL-15/IL-15RA.

The IL-15R is a hetero-trimer composed of IL-15RA, primarily expressed by APCs and the common gamma chain (γ_c_) and IL-2RB, which are expressed by responder T-cells^19^. The γ_c_ family of cytokines (IL-2, -7, -15) all share the γ_c_-receptor subunit but have varying α-subunits^19^. However, cytokine signaling specificity on responder cells is thought to be via the β-subunit^19, 23^. IL-2 and IL-15 share the β-subunit, meaning that the receptor for IL-2 and -15 on responder T-cells is the same (γc and IL-2RB) ^23, 45^. Because of receptor complex is shared it can be difficult to disentangle the relative contributions of IL-2 and -15, though these cytokines have different functions *in vivo*^45^. In turn, this can make it difficult to ascertain the exact function of IL-15 on specific cell types when mixed populations of cells are used in experiments. Thus, IL-15 has been ascribed pro- and anti-inflammatory roles. We attempted to mitigate these concerns by using IL-15/IL-15RA complex and purified CD4^+^ T_RM_ populations from mice and humans, with experiments using APC free *in vitro* systems allowing us to control the presence of IL-15. Thus, one unifying explanation for the discrepant data is that IL-15 has pleiotropic functions that vary by the responder cell type.

Genome-wide association and murine studies have strongly linked T-cells, particularly T_h_17 cells, with the pathogenesis of UC and they are considered major targets of approved therapies like anti-IL-23 agents. Terminally differentiated T-cells are well known to be functionally heterogeneous. Adding to this, T-cells are also anatomically compartmentalized. Thus, naïve CD4^+^ T-cells engage cognate antigens and differentiate into subsets (T_h_17, T_h_1, T_reg_, T_h_2) to eradicate infection (or cause inflammation). Some of these cells are retained long-term and provide systemic (circulating) memory (T_EM_) or reside long-term in peripheral tissues to provide tissue memory (T_RM_). In general, T_EM_ and T_RM_ can be of any subtype of CD4^+^ T-cells, so that circulating subtypes (T_h_17 T_EM_) have tissue resident counterparts (T_h_17 T_RM_)^14, 15^. However, T_RM_ are transcriptionally and functionally distinct from circulating effector memory (T_EM_) T-cells^14, 15^. T_RM_ are highly enriched in tissues suffused with microbes like the intestine, skin, and pulmonary tree, implying they regulate the microbiota.

It has long been held that T_EM_ are the main purveyors of damage in UC. However, T_RM_ are the most abundant T-cell subset in the intestine and are increasingly implicated in the pathogenesis of IBD^7, 14, 15, 17, 18^. Thus, we focused on T_RM_ given that they are abundant, closely aligned with the microbiota (and therefore an excellent candidate to link dysbiotic microbiota with dysregulated host responses), and are increasingly implicated in the pathogenesis of IBD. We specifically focused on CD4^+^ T_RM_ because (CD4^+^) T_h_17 cells are implicated in the pathogenesis of UC and are the target of efficacious therapies for UC. T_RM_ are less accessible than T_EM_ because they are relatively disconnected from the systemic circulation, which makes T_RM_ difficult to target with therapeutic agents. One approach that may circumvent this challenge is to target pathways that regulate the *in situ* biology of T_RM_, hence we focused on IL- 15 which along with the other γ_c_ cytokines regulates homeostatic processes in terminally differentiated cells. IL-15 is predominantly produced in the tissue microenvironment and acts to regulate the local function of resident intestinal CD8^+^ T_RM_ and CD8^+^ intraepithelial lymphocytes (IELs) ^22^. In contrast, the role of IL-15 in CD4^+^ T_RM_ biology is relatively uncharacterized.

Finally, it is critical to recognize that while IL-17 is the defining cytokine of T_h_17 cells, IL- 17(A) is probably not pathogenic in UC. Indeed, two separate clinical trials of anti-IL-17A agents in Crohn’s Disease were stopped early due to poor outcomes in the treatment arm^9^. Consistent with this, IL-17A is protective of the intestinal mucosal in murine models of colitis^10, 11^. In addition, data from human and murine *in vitro* and *in vivo* systems (like experimental autoimmune encephalomyelitis) indicate that pathogenic T_h_17 cells are characterized by the co-production of IL-17A and IFNɣ (‘T_h_1/17’ cells), and that GM-CSF is a major mediator of pathogenic T_h_17 cells^8, 12, 41^. Thus, the prevailing model is that IL-17A is a marker of T_h_17 cells, but that T_h_17 cells are pathogenic via IL-17A-independent pathways. Collectively, our data suggest that IL-15 is upregulated in UC but declines with disease remission, and although IL-15 is redundant for T_h_17 differentiation and the maintenance of T_h_17 T_RM_, it can promote inflammatory T_h_17 T_RM_ via JAK1 pathways.

## METHODS

### Human Subjects

This study conforms with the Helsinki declaration for ethical human subjects’ research and was approved by the University of Michigan Institutional Review Board. All subjects were prospectively recruited and provided written informed consent. Patients with UC and non-IBD controls were identified by screening the IBD and colorectal surgery clinical services. Patients with UC had clinical, histologic, radiographic and laboratory evidence of UC based on standard definitions and were under the care of dedicated IBD specialists. Healthy control subjects included those undergoing surgery for colorectal cancer, diverticulitis, or other benign conditions and did not have any evidence of IBD. Surgical specimens from patients with medically refractory UC and non-IBD controls were obtained via the Tissue Procurement Core of the University of Michigan. All specimens were reviewed by a board-certified GI pathologist to verify inflammation consistent with UC, demarcate areas of quiescence and clear surgical margins, and identify malignancy or infection. GI pathologists had access to all clinical information but were blind to our study groups. Normal control tissue was obtained from the healthy surgical margin of subjects without IBD who were undergoing surgery for colorectal cancer or benign conditions. Subjects with Indeterminate Colitis, Crohn’s Colitis, or gastrointestinal (GI)infection were excluded from this study.

### Mice

Wild-type (WT) C57BL/6 mice were purchased from the Jackson (Bar Harbor, ME) or Taconic (Rensselaer, NY) Laboratories. IL-15^-/-^ mice were obtained from Taconic and are on a C57BL/6 background. This strain has been previously published and detailed information on its derivation can be found on the Taconic website. For all experiments involving IL-15^-/-^ strains, we used Taconic WT C57BL/6 controls. All experiments used 5-8 wk old gender- and age-matched mice, and all mice were housed in specific-pathogen free facilities at the University of Michigan. For all experiments, mice were monitored for health by weight, spontaneous movement, diarrhea, and grooming. This protocol was approved by the University of Michigan Animal Care and Use Committee.

### Golimumab and RISK cohort data

These tissue bulk RNA-sequencing datasets are publicly available from the National Library of Medicine Gene Expression Omnibus (GEO) datasets. The Golimumab clinical trial in UC was accessed using the GSE number GSE92415 and contains sequencing from the rectal mucosa of patients enrolled in this trial. The trial has been published, and Golimumab is approved for use in UC. Bulk sequencing from ileal Crohn’s was obtained from the Pediatric RISK Stratification Study (RISK) cohort, which is a well published cohort of pediatric patients with IBD. This data was accessed as GSE93624.

### Re-analysis of Single-cell RNA-sequencing data

The processed single-cell RNA-transcript counts were collected from three independent studies: (1) the healthy human gut (Gut Cell Atlas; https://www.gutcellatlas.org/, last accessed Nov 14, 2022); (2) inflamed or normal tissue from ulcerative colitis; (3) inflamed or normal tissue from Crohn’s Disease. As each was collected and processed via the same approach (10X Genomics and preprocessed with cell-ranger), we combined data at the per-cell gene count level, in any data format. CD4 and CD8 T-cells were identified by selecting cells with at least one CD4 or CD8 transcript, respectively. The T_RM_ subset was further identified by selecting cells that were CD69^+^, CD45RO^+^, and CCR7^-^. Finally, within the CD4 or CD8 T_RM_ subset, IL2RB^+^ cells were identified. In the subset of CD4 or CD8 T_RM_, we ordinated with UMAP on highly variable genes to confirm the proper integration of the distinct data sources. The proportion of cells in each category was quantified. To identify differentially expressed genes comparing IL2RB^+^ to IL2RB^-^, we used generalized linear modeling (GLM), with the transcript level as the dependent variable and IL2RB^+^ to IL2RB^-^ as the (1 or 0) as the independent variable. A false discovery rate (FDR) was calculated to correct for multiple comparisons with the Benjamini/Hochberg technique. An FDR cutoff of 0.05 was used for significance. The KEGG 2021 Human gene sets were used for enrichment analysis.

### Citrobacter rodentium infection

Wild-type *C. rodentium* (ATCC 51459) from frozen stock was cultured in 5 ml of Luria Bertani (LB) broth supplemented with Ampicillin (100 µg/ml) overnight on a shaker at 200 rpm at 37C to generate viable colonies for infection. A spectrophotometer assessed the colony forming units (CFU)/mL after the overnight incubation. Oral inoculum stocks of 10^9^ CFU/200 µl were prepared by diluting the overnight *C. rodentium* cultures in PBS. Wild-type and IL-15^-/-^ mice were inoculated by oral gavage with 10^9^ CFU in 100 μl. Control mice received an equivalent volume of PBS. To quantify the burden of *C. rodentium*, the stool was weighed and then homogenized in PBS to an even suspension. Serial dilutions of the homogenized stool were plated in triplicate on MacConkey agar plates and incubated for 72 hours at 37°C. Fecal bacterial load was determined by enumerating colonies after the incubation period and was expressed per unit weight of stool (CFU/gm). The bacterial load of mucosal adherent C*. rodentium* was assessed using a piece of colon tissue. The colon was washed with PBS to remove stool and then weighed, homogenized in PBS, and plated in serial dilution.

### Dextran Sodium Sulfate, anti-CD3 enteritis

2.5% DSS solution, prepared fresh, was administered to mice via drinking water for ad libidum intake. Anti-CD3 antibodies (50 µg/20 gm mouse per injection; Bio X Cell, NH) were injected intraperitoneally every other day over 3 days to induce enteritis, as previously published. Weight, grooming, consistency of stool, spontaneous movement, and rectal bleeding were monitored daily.

### Isolation of murine and intestinal and circulating T-cells

Mouse lamina propria mononuclear cells (LPMCs) were isolated with minor modifications to this published protocol. Briefly, the colon was cleared of stool by flushing with cold Hank’s balanced salt solution (HBSS), then washed again with fresh cold HBSS. The tissue was then transferred to a ‘pre-digestion buffer’ composed of warmed HBSS, 2.5% fetal bovine serum (FBS), 5mM EDTA, and 1 mM dithiothreitol, and dissociated using the gentleMACS Dissociator (1 min, Tissue dissociating setting; Mintenyi Biotech, Bregisch Gladbach, Germany). The tissue was then incubated for 20 min at 37C with gentle mixing and was then vortexed (3,000 RPMs, 10s) and washed in fresh cold RMPI-1640. The tissue was transferred to 50 mL conical tubes with 5 mL of digestion solution pre-warmed to 37C and composed of 150 U/ml Collagenase type III (Worthington Biochemical, Freehold, NJ) and 50 µg/ml DNase I, 2% FBS in complete RPMI-1640, and incubated at 37C in a shaker for 30 min (human tissue) or 45 min (mouse tissue). The digested tissue was put through a 100 μm cell strainer, washed in cold HBSS, and centrifuged (450g, 10min, 4C), and the pellet was resuspended in 1 mL of 40% Percoll, and added to a fresh conical with 4 mL of 60% Percoll. The solution was underlayed with 2.5 mL of 80% Percoll to create a 40/80 interface. The samples were centrifuged (860g, 20 min, 21C) and the LPMCs were aspirated from the 40/80 interface. The LPMCs were washed twice with HBSS and resuspended in 1640 RPMI. Human LPMCs were isolated in a slightly modified manner to minimize tissue necrosis and LPMC apoptosis time. Human colon tissue was cleared of fat, washed with cold RPMI, and cut into ∼ 0.5-1 cm pieces. The tissue was then incubated (45 min, 37C) in 50 mL conical tubes and a digestion solution composed of HBSS with Ca+, Mg+, 2% FBS, 0.5 mg/ml DNase I and 0.5 mg/ml Collagenase IV. The resulting cells were strained, washed, and resuspended in complete RPMI. As previously published, human peripheral blood mononuclear cells (PBMCs) were isolated from patients who provided blood samples by Percoll gradient separation. Naive human and mouse T-cells were purified from (human) PBMCs or (mouse) spleen single-cell suspensions and the respective EasySep™ (Vancouver, British Columbia) naive T-cell isolation kits. Cell purity was verified by flow cytometry and was > 95%. CD4^+^ T_RM_ were isolated using flow cytometry-assisted cell sorting (FACS) from LPMCs above by gating on live cells, followed by CD3^+^CD4^+^CD44^+^CD69^+^ (mouse) or CD3^+^CD4^+^CD45RO^+^CD69^+^CCR7^-^ (human) fractions.

### Stimulation of isolated T-cells

For mouse T_h_17 differentiation assays, naive T-cells were plated in 24- or 48-well culture wells for incubation via the T-cell receptor with plate-bound anti-CD3 (10 µg/ml) and soluble anti-CD28 (2µg/ml) with recombinant IL-15/IL-15RA complex (100 ng/ml; Thermo Fischer, Waltham, MA) or recombinant mouse IL-6 (50 ng/ml, BioLegend), TGF- β1 (1ng/mL), IL-23 (5 ng/ml, BioLegend), anti-mouse IL-4 (10 ug/mL, BioLegend), and anti-mouse IFN-γ (10 µg/mL, BioLegend) and incubated for 4 days at 37C. Human T_h_17 differentiation assays were performed similarly except the concentrations of recombinant human IL-6 (30 ng/ml, BioLegend), TGF- β1 (2.25ng/mL), IL-23 (30 ng/ml, BioLegend), anti-human IL-4 (2.5ug/mL), and anti-human IFN-γ (1 µg/mL) and the incubation days (10 days). Purified intestinal murine CD4^+^ T_RM_ were also stimulated with plate-bound anti-CD3 (10 µg/ml) and soluble anti-CD28 (2 µg/ml) with recombinant IL-15/IL-15RA complex (100 ng/ml) or IL-1β (10 ng/ml) and IL-23 (10 ng/ml) and incubated for 4 days in a 37C incubator. Purified human intestinal CD4^+^ T_RM_ were stimulated with plate-bound anti-CD3 (10 µg/ml) and soluble anti- CD28 (2 µg/ml) with recombinant IL-15/IL-15RA complex (100 ng/ml) or IL-1β (50 ng/ml) and IL- 23 (50 ng/ml) and incubated for 4 days in a 37C incubator^46^ . In some experiments, the JAK1 inhibitor filgotinib was also applied (200 nM, Caymen Chemical, Ann Arbor, MI). Media was typically changed every 3 days to maintain cell viability and all supernatants were collected for enzyme-linked immunosorbent assay (ELISA) per kit instructions (R&D Systems, Minneapolis, MN) to quantify cytokine production. Cells were washed and RNA was extracted for qPCR. In some experiments, cells were washed and rested overnight before phorbol 12-myristate 13- acetate and ionomycin stimulated assessment with flow cytometry (FCS).

### Flow cytometry

Cells were rested overnight after purification or stimulation per above in RPMI supplemented with glutamine, sodium pyruvate, 100 units/ml penicillin, 100 µg/ml streptomycin, and 10% fetal bovine serum before stimulation with 1X stimulation cocktail (eBiosciences, Waltham MA) with Golgi Plug (BD Biosciences, Franklin Lakes, NJ) before flow cytometry for 4-6 hours. Flow cytometry was then performed on the LSR II (BD Biosciences, Franklin Lakes, NJ) or the FACS ARIA II (BD Biosciences), and the data were analyzed with FlowJo (Ashland, OR). The following antibodies were used for flow cytometry: CD44 BV 421 (BioLegend), CD103 PE-Dazzle 594 (BioLegend, San Diego, California), IFNγ APC (BioLegend), CD4 FITC (Thermo Fisher, Waltham, MA), Fix Viability eFluor 780 (Thermo Fisher), CD69 PE-Cy7 (BioLegend), IL-17A PE (BioLegend), CD3ε BV 510 (BioLegend), CD8β PerCP-eFluor 710 (eBiosciences), CD62L PE (Miltenyi Biotec, Bergisch Gladbach, Germany), CCR7 APC (Miltenyi Biotec).

### Quantification of cytokines from tissue explants

Mice were euthanized at day 10 after *C. rodentium* oral gavage and the colon (and cecum) were dissected free. The colon was cut longitudinally and washed free of stool. A 2 cm segment of the terminal colon was then cut and placed in a 48-well plate with 400 μl of complete media and incubated at 37C for 24 hours. The supernatant was then collected, and cytokines were measured by ELISA using kit instructions (R&D systems, Minneapolis, MN).

### Quantitative PCR

T-cells or homogenized colon were suspended in RLT buffer (RNAeasy kit, Qiagen, Valencia, CA), RNA was extracted (RNAeasy, Qiagen, Hilden, Germany) and cDNA was synthesized per kit instructions (iScript cDNA Synthesis, Bio-Rad, Hercules, CA). All primers were from Bio-Rad. Some results are also normalized to baseline conditions as the figure legends indicate. Results were analyzed on a CFX Connect system (Bio-Rad).

### Statistics

Experiments with groups of ≥ 3 were analyzed using one-way ANOVA, and experiments with fewer groups were analyzed with paired, or unpaired students *t*-test as appropriate. All data were tested for skewing and normality and parametric or non-parametric tests were applied as appropriate. Most data were analyzed using Prism (GraphPad, San Diego, USA). Due to the complexity of RNA-sequencing data, data in figure 2 as well as predictive modeling were performed separately using R (version 4.2.1).

### Predictive modeling

We again utilized data from the Golimumab clinical trial in UC (GSE number GSE92415) to determine whether *IL15* and related transcript levels from rectal mucosa can predict response to anti-TNF biologic therapies. We performed least absolute shrinkage and selection operator (Lasso) L1-regularized classification model using the R package *glmnet*. Data were randomly split into training (75%) and test set (25%) stratified by the proportion of responders. Lasso models were trained using five-fold cross-validation to estimate the accuracy of the models and to tune hyperparameters. Model accuracy was then evaluated by calculating the area under the receiver operator characteristic curve (AUC) on the independent test set. Variable importance scores were extracted to determine which transcript levels were most important for the model.

### Chemicals

Unless otherwise noted, all chemicals, and media were obtained from Thermo Fisher Scientific (Waltham, MA).

---

**Supplemental Figure 1.**
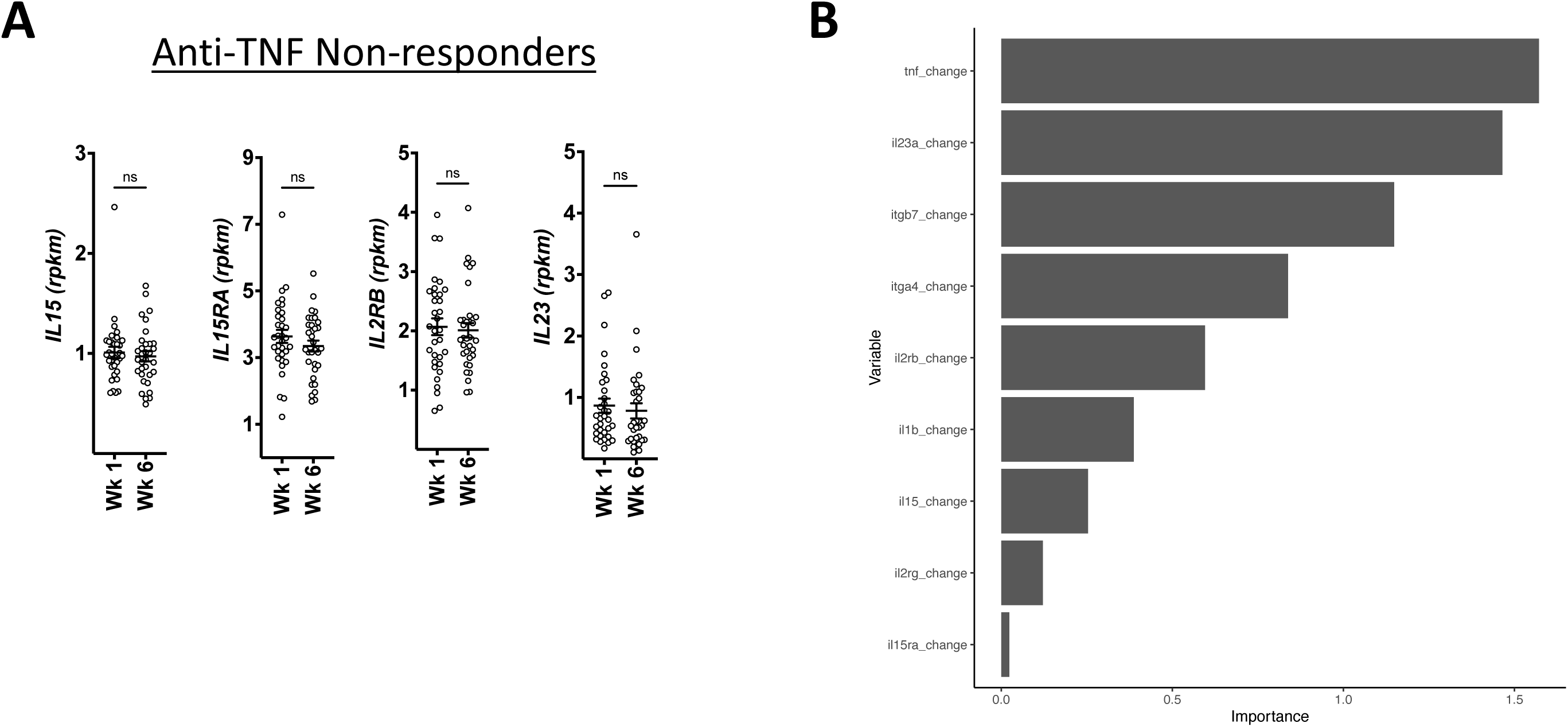
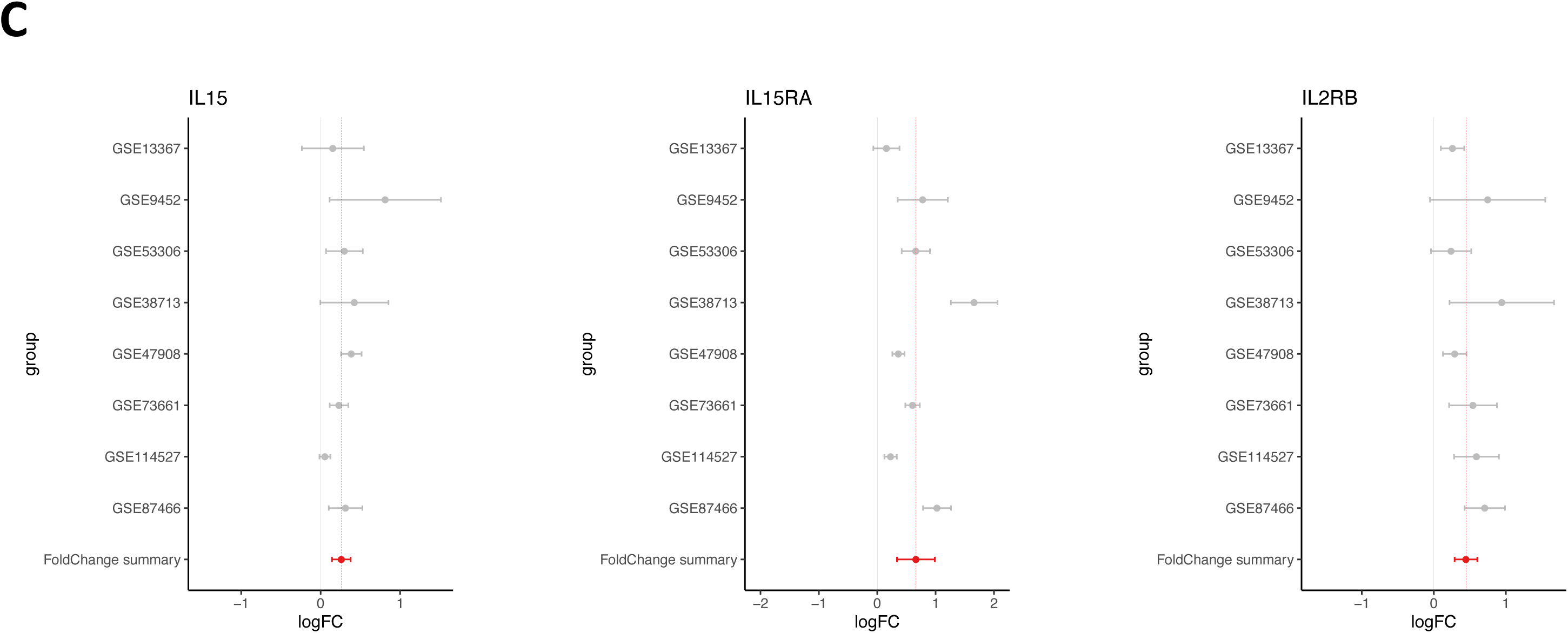
**A)** *IL15* axis transcripts in non-responders to the Golimumab clinical trial. **B)** Predictors of the model used in the least absolute linkage Lasso model to predict response to Goliumuab in Fig 1E, ranked by importance scores. **C)** *IL15* axis transcripts accessed from a meta-analysis of gene transcripts in UC across 85 individual datasets (REF 29).

**Supplemental Figure 2.**
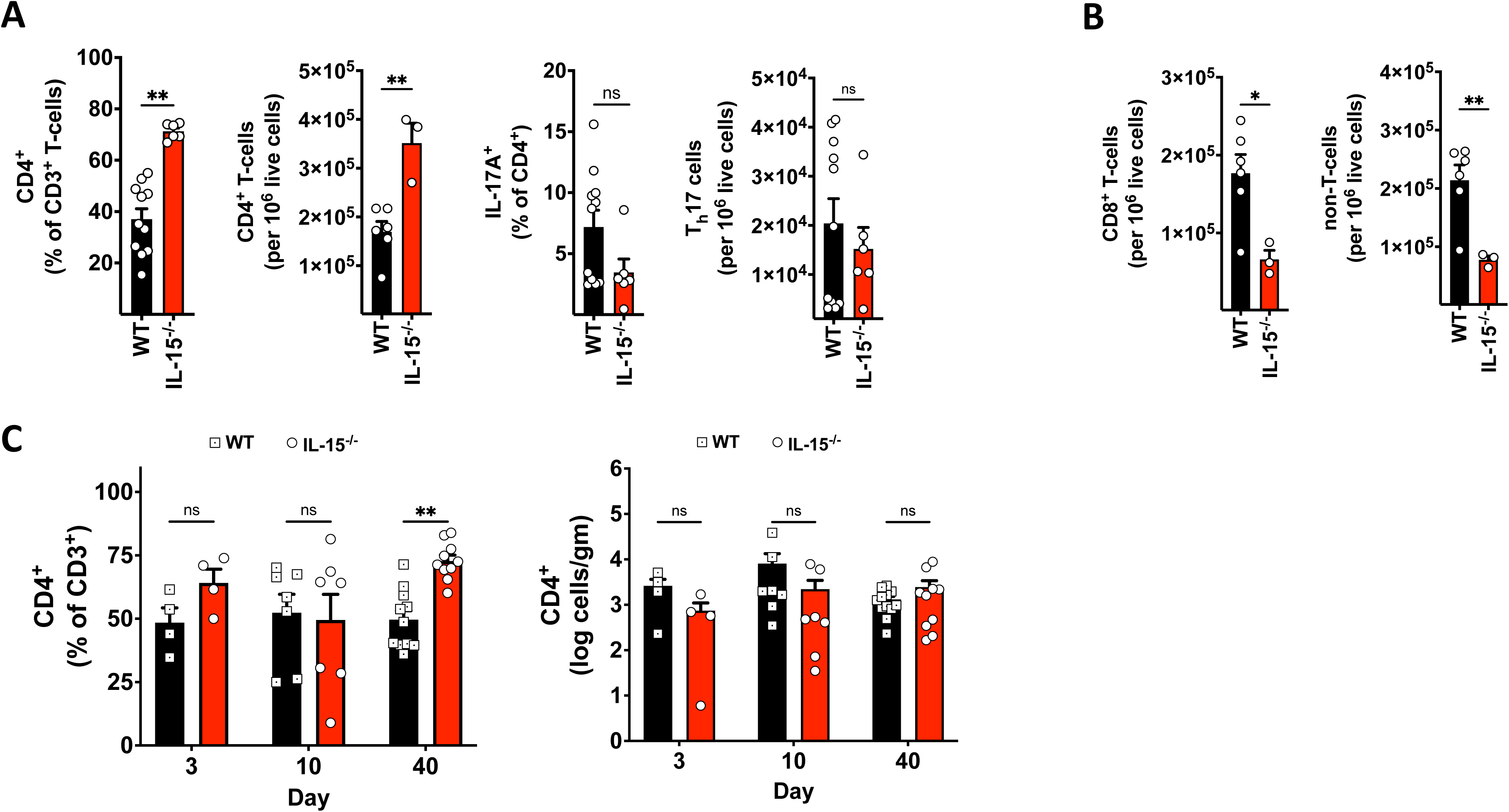
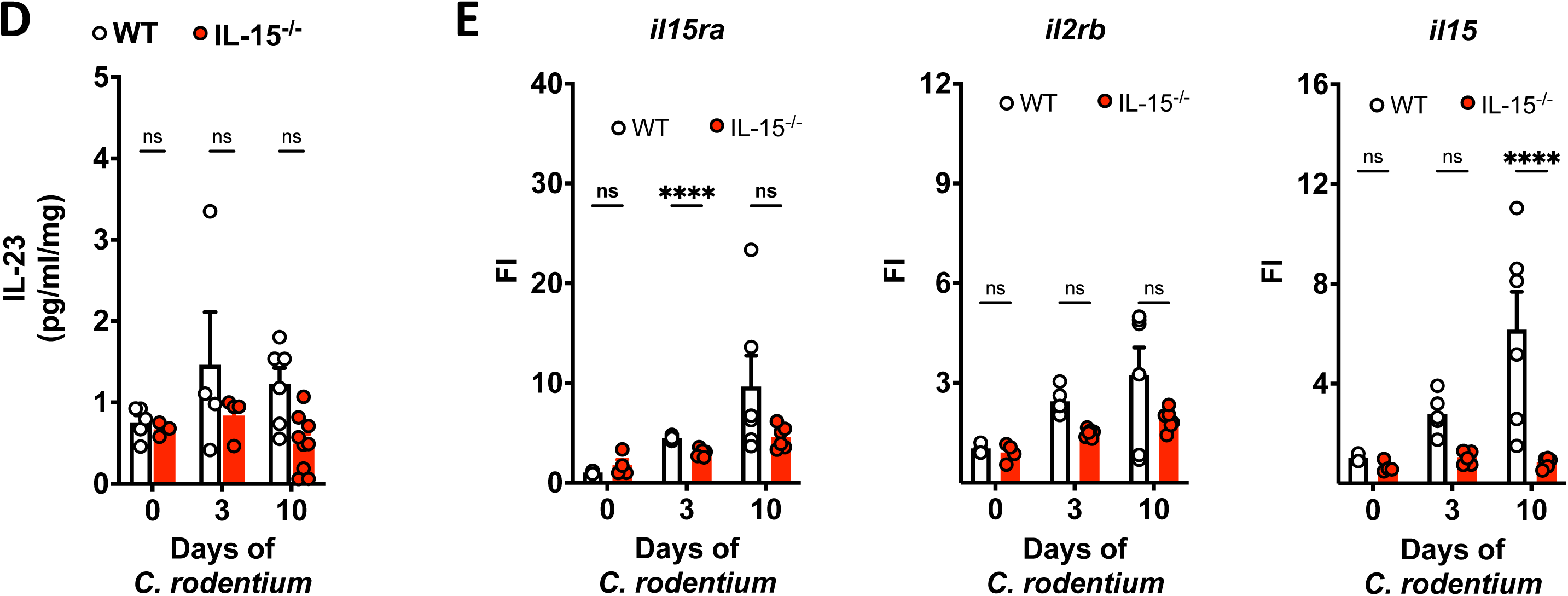
**A)** The percentage and numbers of total CD4^+^ T-cells and T_h_17 cells in the colon of WT and IL-15^-/-^ mice at baseline. **B)** The numbers of total CD8^+^ and non-T-cells in the colon of WT and IL-15^-/-^ mice at baseline. **C)** The percentage (left panel) and total numbers (right panel) of CD4^+^ T-cells in WT and IL-15^-/-^ mice with *C. rodentium*. **D)** IL-23 in colon explants from WT and IL-15^-/-^ mice with *C. rodentium*. **E)** Induction of *il15ra*, *il2rb,* and *il15* in the colon of WT and IL-15^-/-^ mice with *C. rodentium*.

**Supplemental Figure 3.**
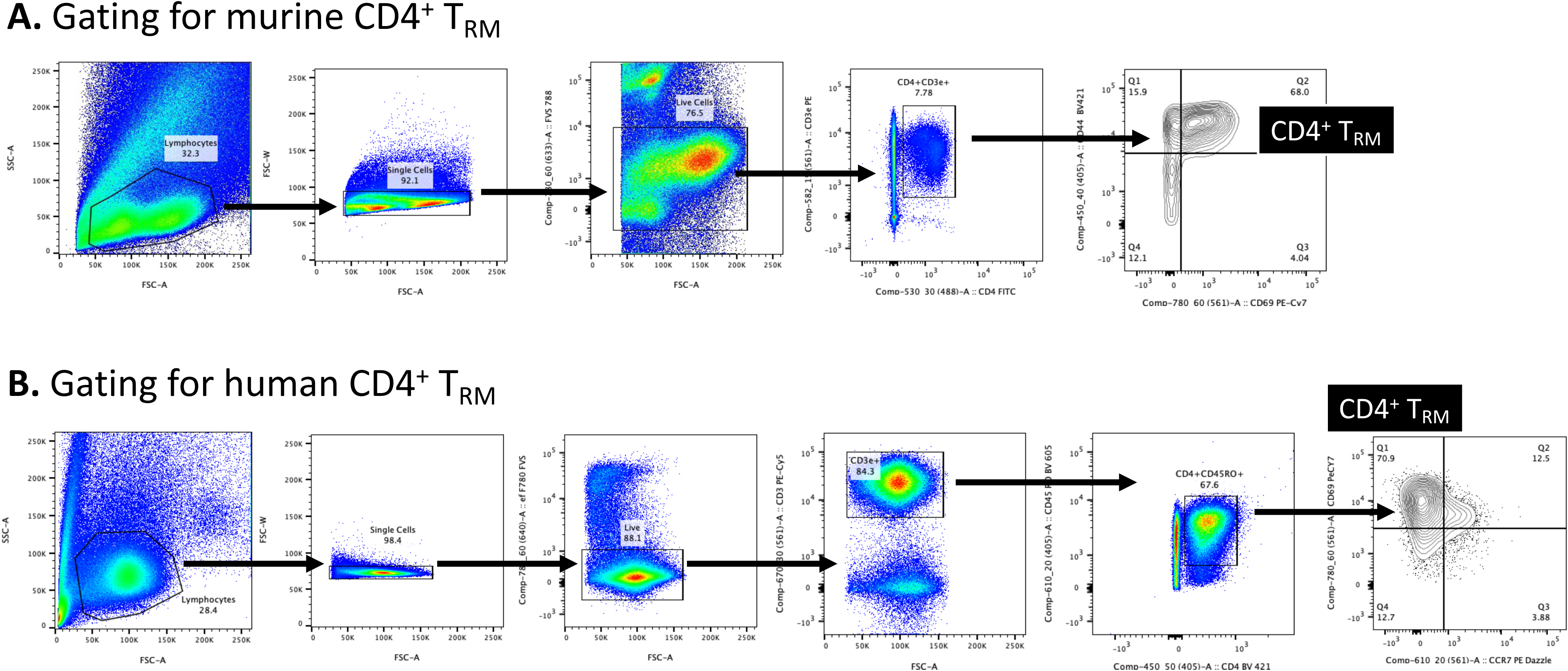
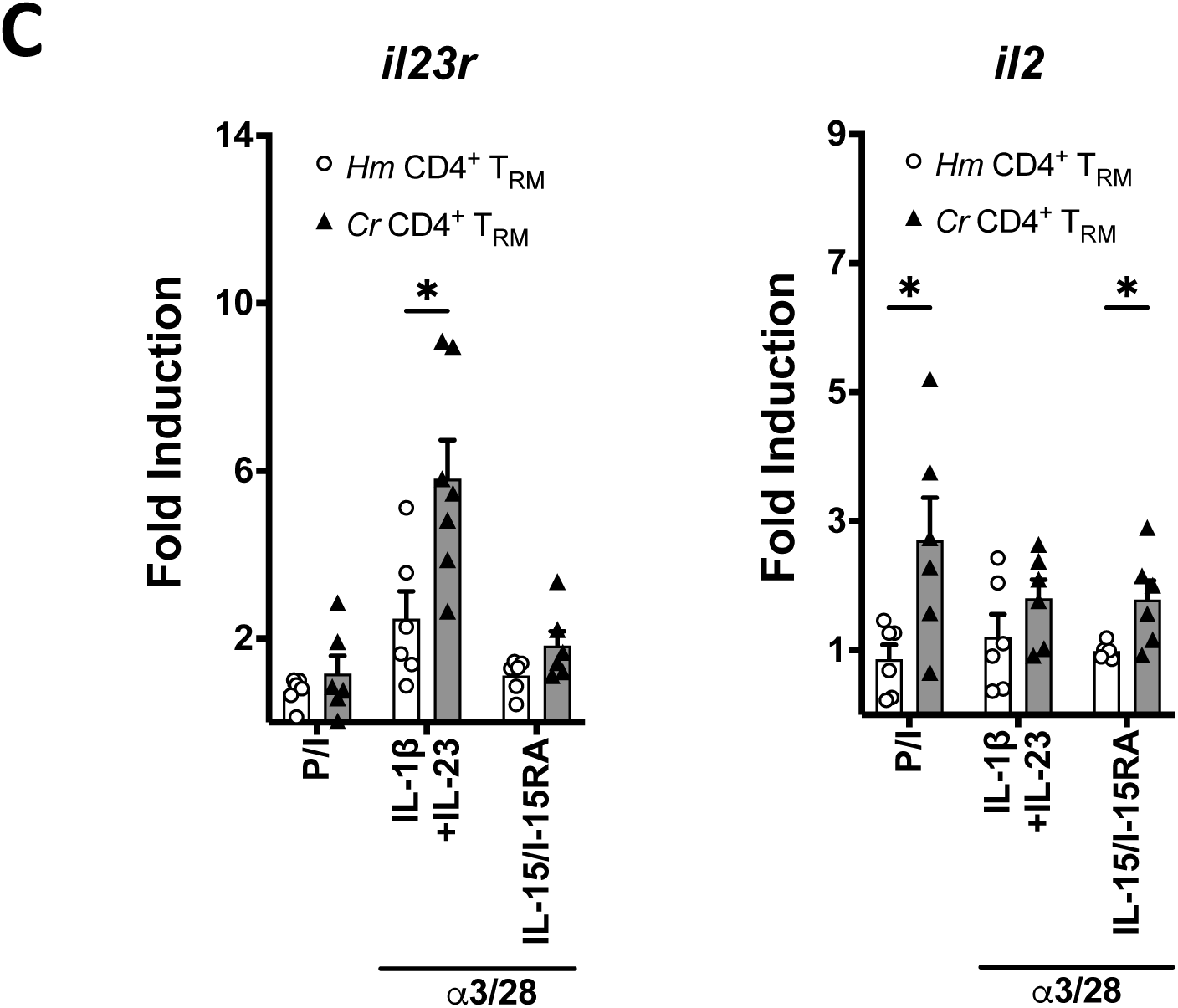
**A)** FACS purification of colonic CD4^+^ T_RM_ from **(A)** mice and **(B)** humans as in Figs 5A and C, respectively. **C)** Colonic CD4^+^ T_RM_ were isolated and stimulated as in Fig 5A and the induction of *il23r* and *il2* was determined. Genes and conditions are normalized to GAPDH and *Hm* CD4^+^ T_RM_, respectively. * = p< 0.1

